# High similarity among ChEC-seq datasets

**DOI:** 10.1101/2021.02.04.429774

**Authors:** Chitvan Mittal, Matthew J. Rossi, B. Franklin Pugh

## Abstract

ChEC-seq is a method used to identify protein-DNA interactions across a genome. It involves fusing micrococcal nuclease (MNase) to a protein of interest. In principle, specific genome-wide interactions of the fusion protein with chromatin result in local DNA cleavages that can be mapped by DNA sequencing. ChEC-seq has been used to draw conclusions about broad gene-specificities of certain protein-DNA interactions. In particular, the transcriptional regulators SAGA, TFIID, and Mediator are reported to generally occupy the promoter/UAS of genes transcribed by RNA polymerase II in yeast. Here we compare published yeast ChEC-seq data performed with a variety of protein fusions across essentially all genes, and find high similarities with negative controls. We conclude that ChEC-seq patterning for SAGA, TFIID, and Mediator differ little from background at most promoter regions, and thus cannot be used to draw conclusions about broad gene specificity of these factors.

---

ChEC-seq (Chromatin Endonuclease Cleavage-sequencing) is a method aimed at tracking protein-DNA interactions across a genome (Zentner et al., 2015). The concept of ChEC-seq is to fuse the DNA coding sequence of micrococcal nuclease (MNase) to the coding sequence of a chromatin-associated protein. Genome-wide binding of these MNase fusion proteins are considered innocuous to DNA cleavage until cells are made permeable to calcium ions, wherein the MNase is activated. Activated MNase cleaves the local DNA regions to which the fusion protein is bound, thereby providing a genomic mark of binding. Cleavages can be quantified by the frequency of DNA ends detected by deep sequencing. ChEC-seq has been applied to several chromatin organizing proteins including the site-specific General Regulatory Factors (GRFs) called Abf1, Reb1, and Rap1 (Zentner et al., 2015). ChEC-seq has also been utilized to assess binding profiles of Mediator, SAGA, TFIID, and RSC chromatin remodeler complexes, all of which play critical roles in transcription initiation by RNA polymerase II (Baptista et al., 2017; Donczew et al., 2020; Grunberg et al., 2016). It has also been applied to Rif1, which predominantly binds telomeres rather than promoter regions (Hafner et al., 2018), and thus potentially serves as an independent negative control at promoters.

Interpretation of ChEC-seq data has led to conclusions that contrast with prior studies. In one, it was reported that GRFs bind a subset of sites based primarily on DNA shape rather than on direct sequence readout (Zentner et al., 2015). However, a follow-up re-analysis of the data challenged the validity of such conclusions based on ChEC-seq (Rossi et al., 2017; Zentner et al., 2017). We therefore re-examined the published yeast ChEC-seq data of SAGA, TFIID, and Mediator subunits. For comparison, we also included RSC, Rif1 and GRF ChEC-seq data, and negative controls that essentially employed unfused MNase. Cleavage sites were examined around transcription start sites of all 5,716 previously-defined yeast coding genes (Xu et al., 2009) (rows in **Figure 1**). Genes were segregated into three previously-defined classes (Basehoar et al., 2004; Huisinga and Pugh, 2004; Reja et al., 2015): Ribosomal protein (RP) genes, SAGA-dominated genes, and TFIID-dominated genes. As in prior studies (Baptista et al., 2017; Grunberg et al., 2016; Zentner et al., 2015), ChEC-seq data (DNA cleavages) were globally normalized such that the total number of sequencing reads being plotted in each dataset were the same.

**Figure 1.**
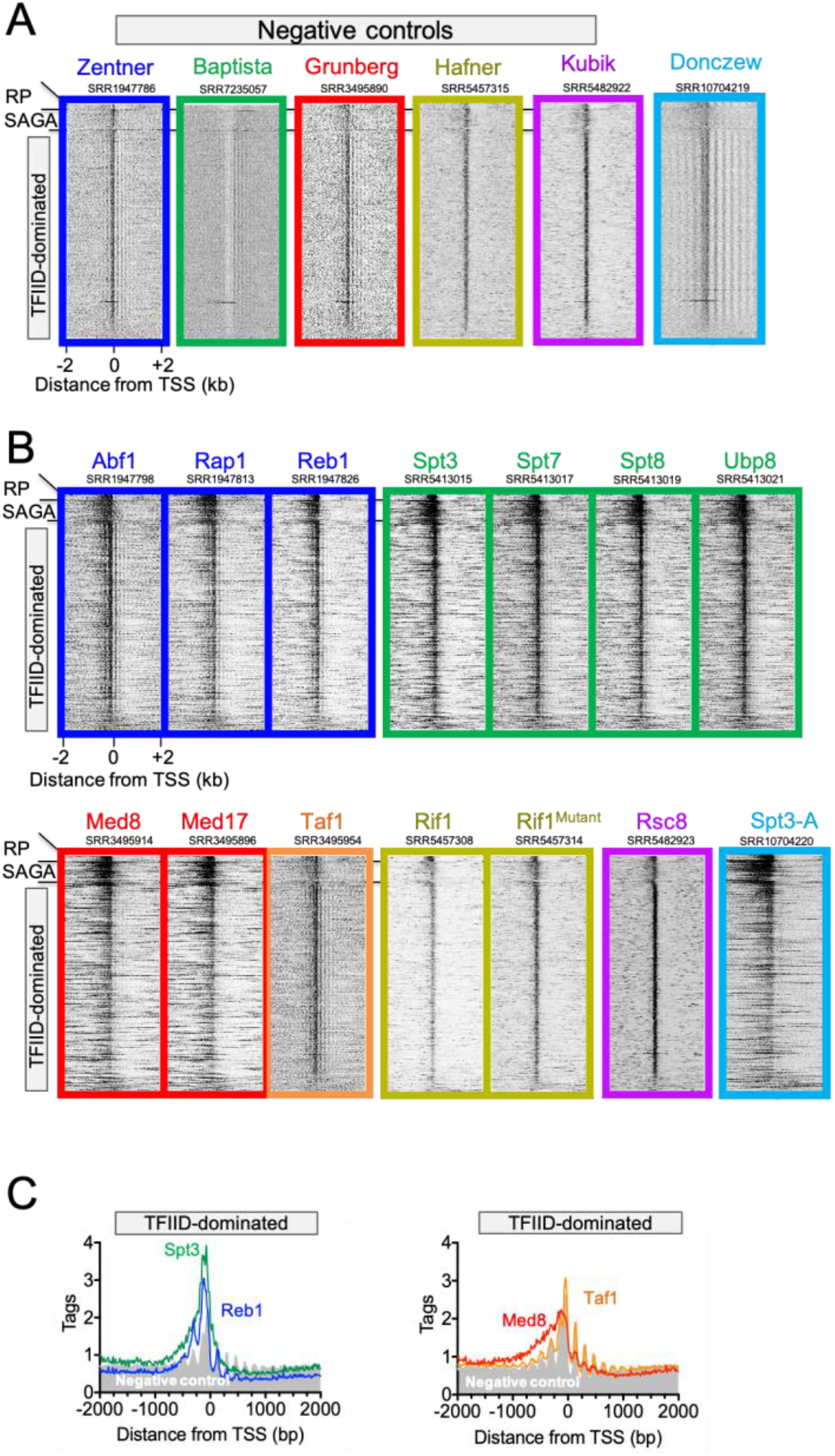
Plots of published ChEC-seq data at 5,716 yeast genes. **A-B**, Heatmaps of ChEC-seq negative controls (**A**) or test samples (**B**) in which genes were separated into the indicated three classes (RP denotes Ribosomal Protein genes, SAGA denotes SAGA-dominated genes) (Huisinga and Pugh, 2004), then sorted by transcript level (high to low) (Yassour et al., 2009) within each class. Frames of similar color are from the same study (indicated above the negative controls along with accession numbers). Analyses were conducted on datasets derived from 5 minutes of MNase cleavage, which was the time point used in the SAGA/TFIID/Mediator studies. Some studies did not report or provide datasets for a matched negative control. We therefore used whichever control dataset was available: Baptista (15 min negative control), and Kubik (20 min. negative control, 10 sec. Rsc8). **C**, Average MNase cleavage (tag counts) for TFIID-dominated genes for selected datasets from panels A-B.

We first compared the available ChEC-seq negative controls (unfused MNase) across each study (**Figure 1A**). Although not tested or reported in prior studies, an implicit assumption is that the protein level of the unfused MNase control and its localization to the nucleus are similar to each fusion protein being tested. This presents a particular concern where local background cleavages are similar in magnitude as in test samples, and insufficient background replicates are available to assess variability. With one exception (Baptista), the negative controls from each study had widespread enrichment of MNase cleavages in nucleosome-depleted or nucleosome-free promoter regions (NDR/NFR). In addition, cleavages existed in the linker regions between nucleosomes. Both types of cleavages reflect nonspecific background by MNase that is expected of all MNase fusion proteins. These cleavages are due to higher accessibility of linker DNA to MNase, where histones/nucleosomes are absent or depleted. Cleavages in NDRs are expected to be more frequent than in linkers because NDRs contain ten times more accessible DNA (~150 bp vs ~15 bp). The Baptista negative control was an outlier in our analysis, and thus was not analyzed further. It appeared to be over-digested, perhaps due to a reported 15-minute MNase digestion instead of 5 minutes as used for the test samples. This gives the appearance of low background, but instead may be due to small DNA fragments not forming mappable libraries. See Methods for further information on this.

One key observation stood out with nearly all ChEC-seq datasets: cleavage patterns across all genes looked strikingly similar when different datasets were compared to each other and with the negative controls (globally in **Figure 1B**). Similarities were most consistent at the TFIID-dominated (n=5,081) classes of genes, representing ~90% of all genes. When quantified and averaged (**Figure 1C**), little difference from the negative controls was observed (see gray-filled plots compared to colored traces), indicating modest or no factor enrichment over background. For example, Taf1 at TFIID-dominated genes showed essentially no enrichment over the reported negative control (orange trace vs. gray fill in **Figure 1C**). Note that Rif1 and its DNA-binding mutant are not expected, nor observed, to be enriched at promoter regions compared to controls. Yet, such regions are intrinsically hyper-sensitive to MNase relative to gene bodies. We note that higher site-specificity for GRFs was reported in the Zentner study when using shorter digestion times up to ~1 minute, instead of the 5 minutes examined here. In those studies, sequence-specific binding was detectable above background. For the purposes of examining SAGA, TFIID, and Mediator, we analyzed the 5-minute timepoints because it was the time point from which conclusions in those papers were drawn, and the only time point that was present consistently across all studies.

One noticeable difference was at the 501 SAGA-dominated genes (“SAGA” in **Figure 1A,B**). SAGA, Mediator, and GRF ChEC-seq data, while similar to each other, were distinctly enriched relative to their respective negative controls and from TFIID (Taf1), RSC (Rsc8) and Rif1. Taken at face value, such patterning might indicate that SAGA, Mediator, and GRFs, but not TFIID or RSC, are predominantly at SAGA-dominated genes. Such tentative conclusions on SAGA and TFIID run counter to the conclusions from the published ChEC-seq studies (Baptista et al., 2017; Grunberg et al., 2016; Warfield et al., 2017), where it was reported that SAGA, TFIID, and Mediator are present at most genes in both classes. However, further inspection (below) of the data led us to question any interpretation of ChEC-seq data at even SAGA-dominated genes.

Since SAGA, TFIID, and Mediator do not bind specific DNA sequences, we could not independently verify their binding through DNA motif enrichment. However, GRFs have cognate motifs. Thus, cognate motif enrichment should coincide with the observed GRF ChEC-seq enrichment at SAGA-dominated genes and serve as a positive control for SAGA, TFIID, and Mediator binding at these genes. Surprisingly, we found little or no enrichment of GRF motifs at SAGA-dominated genes, despite ChEC-seq enrichment (**Figure 2**, left panels). For comparison, Reb1 and Abf1 motifs were enriched at TFIID dominated genes (**Figure 2**, right panels). Therefore, the GRF ChEC-seq enrichment that occurs at SAGA-dominated genes was not supported by cognate motif enrichment, and thus was not independently validated. While we do not exclude the possibility that GRF enrichment at SAGA-dominated genes occurs by some indirect mechanism, taken at face value we find no independent validation of the ChEC-seq assay as applied to SAGA, TFIID, and Mediator.

**Figure 2.**
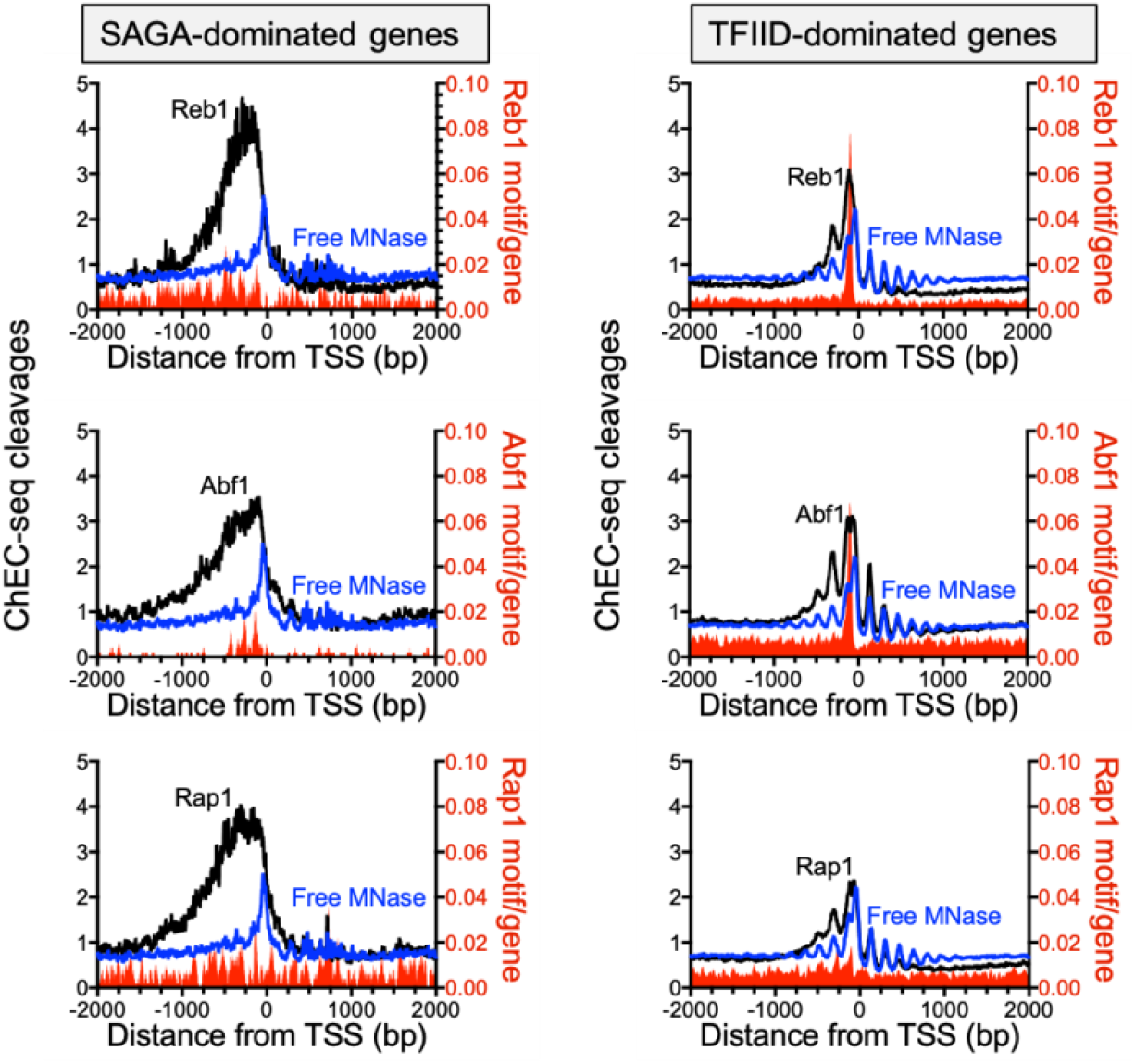
ChEC-seq data for GRFs at SAGA-dominated genes are not validated by motif occurrence. Average cleavage frequencies for Reb1, Abf1 and Rap1 ChEC-seq data (black) or free-MNase negative control (blue) were plotted at the SAGA-dominated (left panel) or the TFIID-dominated (right panel) gene classes (Huisinga and Pugh, 2004). Normalized motif occurrences for each GRF are also plotted (red fill).

One of the surprising conclusions from the ChEC-seq studies on SAGA, TFIID, Mediator, and GRFs is their general presence at most genes, much as one would expect for a general transcription initiation factor. To address this at a quantitative level, we segregated genes into those that were most enriched for a factor’s ChEC-seq signal (top 25%), and the rest (bottom 75%). As shown in **Figure 3**, we observed no ChEC-seq enrichment relative to free MNase for the bottom 75%, regardless of the tested factor. Thus, the ChEC-seq data do not support the conclusion that these factors are at most genes.

**Figure 3.**
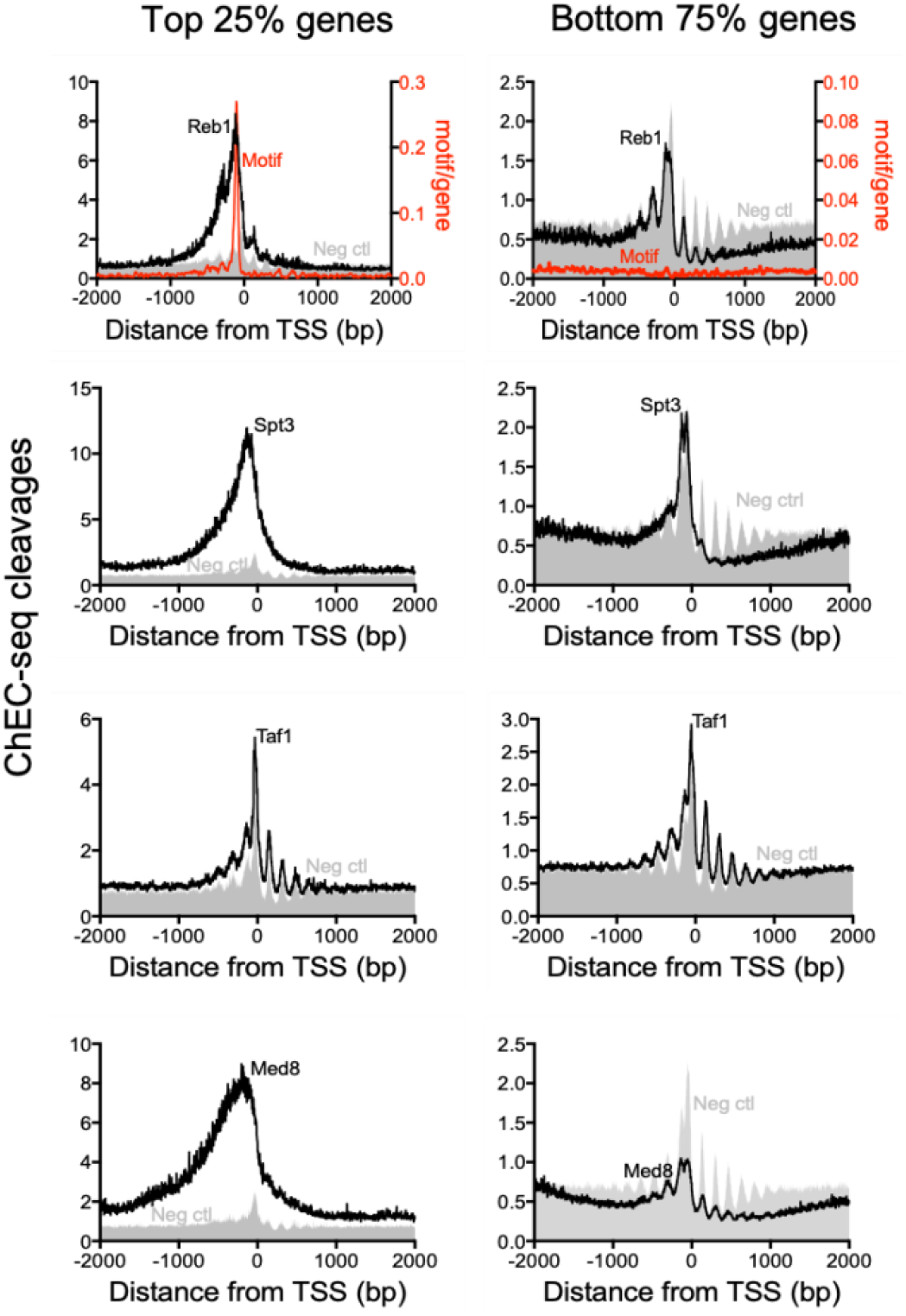
ChEC-seq cleavages are not enriched at most genes. Datasets were the same as from Figure 1. All genes were sorted based on cleavage frequency of the indicated factor, then averaged for the top 25% (left panels) and the bottom 75% (right panels). Averages for the negative control (Zentner dataset in all cases) was for the same sets of genes. Note differences in Y-axis values in left vs right plots. For Reb1 plots (upper panels), motif occurrence (red trace) is also shown, which validates Reb1 specificity at sites of high cleavage.

In summary, we conclude that the ChEC-seq assay, as implemented and/or analyzed in prior studies on SAGA, TFIID, Mediator, and GRFs, lacks sufficient specificity to support conclusions regarding the proposed broad gene specificity of these factors.

## Methods

### Analysis of ChEC-seq data

Raw FastQC files for the ChEC-seq datasets were downloaded from GEO using the dataset accession number indicated above each figure panel. We note that the Baptista negative control data file (pSpt3_free_MNase_15min SRR7235057, from SRX4141422 in GSM3165083) reflects a later upload from May 30, 2018 as reported in a Correction (Baptista et al., 2018). The 5-minute negative control dataset was not available. We therefore used the 5-minute negative controls from the Zentner and Gruenberg studies. Files were mapped to SacCer3 genome using BWA to generate BAM files. Datasets were normalized such that total tags in each dataset were set to be equal, in accord with methods accompanying the published datasets (Baptista et al., 2017; Grunberg et al., 2016; Zentner et al., 2015). Normalized data were mapped to the TSSs of 5,716 genes transcribed by RNA polymerase II (Xu et al., 2009).

### Data analysis

Analyses were performed using Scriptmanager v.011, which is publicly available for download at: https://github.com/CEGRcode/scriptmanager. Reference files used to map datasets in this study can be found at: https://github.com/CEGRcode/Mittal_2021_ChEC-seq.

### Heatmaps and composite plots for Figure 1

Tag Pileup function of Scriptmanager v0.11 was used to generate the heatmaps and composite plots using the following settings: Read_1, Strands combined, 0 bp tag shift, 1 bp bin size, set tags to be equal, sliding window 3. Equivalent results were obtained with larger bin sizes (tested up to 25 bp). Output files for heatmaps were CDT files, which were visualized in Java Treeview (fixed contrast 3.0). Output files also included Composite_Average.out files, which were plotted using Prism 7 software to generate composite plots.

### Composite plots for Figure 2

Datasets were mapped to transcription start sites (TSS) of SAGA-dominated and TFIID-dominated genes (Huisinga and Pugh, 2004) (Yassour et al., 2009), using the Tag Pileup function of Scriptmanager v0.11, as described for Fig. 1, except the sliding window was 9. Expanded FIMO BED files were mapped to the abovementioned TSS BED files (expanded to 4 kb) using the Align BED to Reference feature in Scriptmanager. Motif occurrences were summed across a 4 kb window from the TSS and were then divided by number of genes in each class to get average motif occurrence, across the 4 kb window. ChEC-seq cleavages and motif occurrence per gene were plotted together on Prism 7.

### Composite plots for Figure 3

Datasets were mapped to TSSs of all 5716 genes, using the same parameters as in Figure 1. ChEC-seq cleavages were summed across a 500 bp window centered at the TSS, and sorted from highest to lowest. TSS-centric BED files were generated from the upper 25% and lower 75%, along with a free MNase control (Zentner) for the same set of genes.

## Acknowledgements

This work was supported by National Institutes of Health (NIH) grant ES013768 (B.F.P.).

